# The Soybean Expression Atlas v2: a comprehensive database of over 5000 RNA-seq samples

**DOI:** 10.1101/2023.04.28.538661

**Authors:** Fabricio Almeida-Silva, Francisnei Pedrosa-Silva, Thiago M. Venancio

**Affiliations:** Department of Plant Biotechnology and Bioinformatics, Ghent University, 9052 Ghent, Belgium; VIB Center for Plant Systems Biology, VIB, 9052 Ghent, Belgium; Laboratório de Química e Função de Proteínas e Peptídeos, Centro de Biociências e Biotecnologia, Universidade Estadual do Norte Fluminense Darcy Ribeiro, Campos dos Goytacazes, Brazil

**Keywords:** transcriptomics, bioinformatics, gene expression, integrative biology

## Abstract

Soybean is a crucial crop worldwide, used as a source of food, feed, and industrial products due to its high protein and oil content. Previously, the rapid accumulation of soybean RNA-seq data in public databases and the computational challenges of processing raw RNA-seq data motivated us to develop the Soybean Expression Atlas, a gene expression database of over a thousand RNA-seq samples. Over the past few years, our database has allowed researchers to explore the expression profiles of important gene families, discover genes associated with agronomic traits, and understand the transcriptional dynamic of cellular processes. Here, we present the Soybean Expression Atlas v2, an updated version of our database with a 4-fold increase in the number of samples, featuring transcript- and gene-level transcript abundance matrices for 5481 publicly available RNA-seq samples. New features in our database include the availability of transcript-level abundance estimates and equivalence classes to explore differential transcript usage, abundance estimates in bias-corrected counts to increase the accuracy of differential gene expression analyses, a new web interface with improved data visualization and user experience, and a reproducible and scalable pipeline available as an R package. The Soybean Expression Atlas v2 is available at https://soyatlas.venanciogroup.uenf.br/, and it will accelerate soybean research, empowering researchers with high-quality and easily accessible gene expression data.

## Introduction

Soybean (*Glycine max* [L.] Merr.) is one of the most important crops worldwide due to its high protein and oil content, and it is used as a source of food, feed, and industrial products. The availability of its genome sequence (Schmutz *et al*., 2010) and advances in RNA sequencing have revolutionized our understanding of the molecular mechanisms underlying important traits, such as yield (Lu *et al*., 2021; Yang *et al*., 2021), stress tolerance (Tamang *et al*., 2021; Wang *et al*., 2021), and seed composition (Jones and Vodkin, 2013; Lu *et al*., 2016). RNA-seq data of soybean are accumulating at an unprecedented rate in public databases, enabling researchers to access a vast amount of transcriptomic information for gene expression analysis and functional studies (Almeida-Silva *et al*., 2021). However, obtaining gene-level transcript abundance (*i*.*e*., “gene expression”) matrices from raw RNA-seq data is time consuming and requires heavy computational resources.

To address this issue, we previously developed the Soybean Expression Atlas, a gene expression database comprising 1298 publicly available RNA-seq samples that were systematically collected, processed, and quantified using a unified pipeline (Machado *et al*., 2020). Since its release, the Soybean Expression Atlas has allowed researchers to explore the expression profiles of important gene families (Almeida-Silva and Venancio, 2022; Coelho *et al*., 2022; Hou *et al*., 2022; Huang *et al*., 2021; Sangi *et al*., 2021), discover genes associated with agronomic traits (Zhang *et al*., 2022; Almeida-Silva and Venancio, 2021; Almeida-Silva and Venancio, 2023a; Turquetti-Moraes *et al*., 2022; Guo *et al*., 2022), and understand the dynamics of gene regulation underlying plant physiology (Almeida-Silva *et al*., 2020; Sangi *et al*., 2023; Chen *et al*., 2023; Li *et al*., 2021). Up to April 2023, our database has been accessed by more than ten thousand users, 60% of which are recurring. However, a vast amount of new RNA-seq samples have been deposited in public databases since then, creating an urge for an updated version of our database.

Here, we present the Soybean Expression v2 (https://soyatlas.venanciogroup.uenf.br/), an updated version of our database comprising transcript- and gene-level transcript abundance matrices for 5481 publicly available RNA-seq samples. Besides the 4-fold increase in number of samples, the new features of the Soybean Expression Atlas v2 include: i. the availability of transcript-level abundance estimates and equivalence classes, which can be used to explore differential transcript usage; ii. abundance estimates in bias-corrected counts, which corrects for technical biases in raw read counts, increasing the accuracy of differential gene expression analyses; iii. a new web interface with significant improvements in data visualization and user experience, and; iv. a reproducible and scalable pipeline available as an R package. The Soybean Expression Atlas v2 will keep accelerating soybean research, empowering researchers with high-quality and easily accessible gene expression data.

## Materials and Methods

### Reference genome data acquisition

All genomic data used in this version of the Soybean Expression Atlas were based on the Wm82.a4.v1 assembly of the *Glycine max* genome. The reference transcriptome and transcript-to-gene ID mappings were downloaded from PLAZA Dicots 5.0 (Van Bel *et al*., 2022).

### Obtaining transcript abundances from raw sequencing data

To make our pipeline reproducible, we developed an R package named *bears* (https://github.com/almeidasilvaf/bears) that integrates multiple RNA-seq software tools, allowing users to estimate transcript- and gene-level transcript abundances from raw RNA sequencing data available on NCBI’s Sequence Read Archive (SRA) (Sayers *et al*., 2023). The package supports transcript abundance estimation using both alignment-based approaches (i.e., consisting of aligning reads to a reference genome followed by quantification) and lightweight mapping approaches (i.e., direct abundance estimation through pseudomapping or selective alignment to a reference transcriptome). In this version of the Soybean Expression Atlas, we only used the lightweight mapping approach, as it is dramatically faster than alignment-based approaches while also accounting for technical biases affecting transcript quantification (‘t Hoen *et al*., 2013; Patro *et al*., 2017).

Metadata for all RNA-seq experiments available on the SRA (up to June 2022) were obtained with the function *create_sample_info()* and filtered to remove experiments using SOLiD sequencing (due to its obsolescence) and long-read technologies (which are typically used for transcript assembly, not for quantification). FASTQ files were downloaded from the European Nucleotide Archive’s mirror of the SRA using the function *download_from_ena()*, and file integrity was checked with the *check_md5()* function. Adapters and low-quality bases were identified and removed with fastp (Chen *et al*., 2018) using the function *trim_reads()*. FASTQ files with mean read length <40 and/or Q20 rate <80% after filtering were considered as having insufficient quality and removed. Transcript abundances for the samples that passed the sequence quality check were estimated with salmon (Patro *et al*., 2017) using the function *salmon_quantify()*, which performs mapping to a reference transcriptome followed by quantification. Finally, we removed samples for which <50% of the reads failed to map. Salmon was run with the ‘--dumpEq’ flag to generate equivalence classes that can be used in differential transcript usage analyses.

### Summarizing transcript-level to gene-level abundances

Gene-level transcript abundances were obtained with the function *salmon2se()*, which reads quant.sf files created by salmon and creates a *SummarizedExperiment* object containing gene-level abundances in transcripts per million (TPM) and read counts. The count matrix is corrected for biases using the “bias correction without an offset” method implemented in the tximport package (Soneson *et al*., 2015), which performs scaling using the average transcript length over samples and library size.

### Dimensionality reduction

To maximize biological signal and reduce noise, we only used genes with highly variable transcriptional patterns for dimensionality reduction. We used the scran package (Lun *et al*., 2016) to model the mean-variance relationship in a matrix of log-transformed counts normalized by library size, followed by the extraction of the top 5000 genes with the highest biological components (Supplementary Figure S1A). We performed a principal component analysis (PCA) of the highly variable genes and extracted the top 8 principal components to perform dimensionality reduction with *t-*stochastic neighbor embedding (t-SNE) and uniform manifold approximation and projection (UMAP) (Supplementary Figure S1B). As t-SNE and UMAP results can vary depending on perplexity values and number of neighbors, respectively, we ran both algorithms with 6 different values (10, 20, 30, 40, 50, and 60) and selected the optimal value based on visual inspection. The optimal perplexity value for the t-SNE was 60, and the optimal number of neighbors for the UMAP was 30.

### Construction of the web application

The web application is a Shiny app that uses the shinydashboard template (Chang *et al*., 2021; Chang and Ribeiro, 2019). The gene expression database used in the app is stored in a partitioned parquet directory that can be downloaded from the FigShare repository associated with this manuscript (Almeida-Silva and Venancio, 2023b). The interface between R and the Apache Arrow platform is performed with the arrow R package (Richardson *et al*., 2021). Data visualization is powered by ggplot2 (Wicham, 2016), ComplexHeatmap (Gu, 2022), and plotly (Plotly Technologies Inc, 2015).

## Results and Discussion

### The Soybean Expression Atlas comprises 5481 high-quality samples

We downloaded 6921 RNA-seq samples available on the European Nucleotide Archive (ENA) and processed them in a unified pipeline to generate a large database of gene-level transcript abundances (Figure 1A, see Materials and Methods for details). After trimming reads to remove adapters and low-quality bases, we performed the first quality check, which consists in removing samples with mean read length <40 bp and/or Q20 rate <80%. 158 samples did not pass this quality check and were removed (Figure 1B, Supplementary Table S1). Further, after mapping reads to the reference transcriptome, we performed the second checkpoint, consisting of removing samples with mapping rates <50%. This step led to the removal of 1282 samples (Figure 2B, Supplementary Table S2), resulting in the final Soybean Expression Atlas v2 with 5481 samples (Supplementary Table S3).

**Fig. 1.**
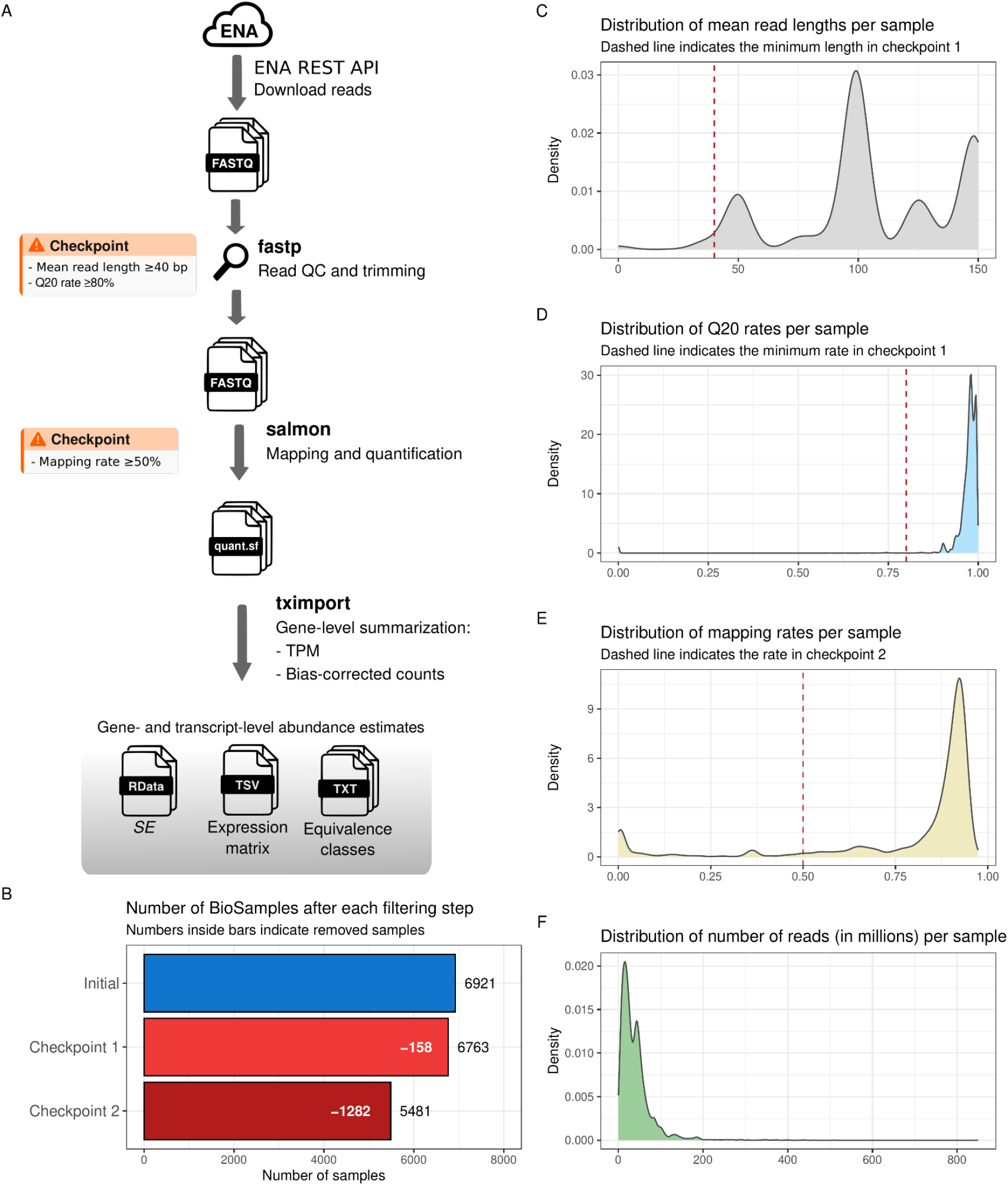
Pipeline used to create the Soybean Expression Atlas v2 and summary quality statistics. **A**. Data acquisition and processing pipeline used in the Soybean Expression Atlas v2 (see Materials and Methods for details). Checkpoints represent stages where samples were removed. All data generated at the end of the pipeline (*SummarizedExperiment* objects for gene- and transcript-level transcript abundances and equivalence classes) are available in the FigShare repository associated with this publication. **B**. Number of BioSamples before and after each filtering step. Numbers next to the bars represent the remaining samples at each step, and numbers inside bars represent removed samples relative to the previous step. **C**. Distribution of mean read lengths per sample. Dashed red line represents the minimum read length threshold from checkpoint 1. **D**. Distribution of Q20 rates per sample. Dashed red line represents the minimum Q20 rate threshold from checkpoint 1. **E**. Distribution of mapping rates per sample. Dashed red line indicates the minimum mapping rate threshold from checkpoint 2. **F**. Distribution of number of reads (in millions) per sample.

**Fig. 2.**
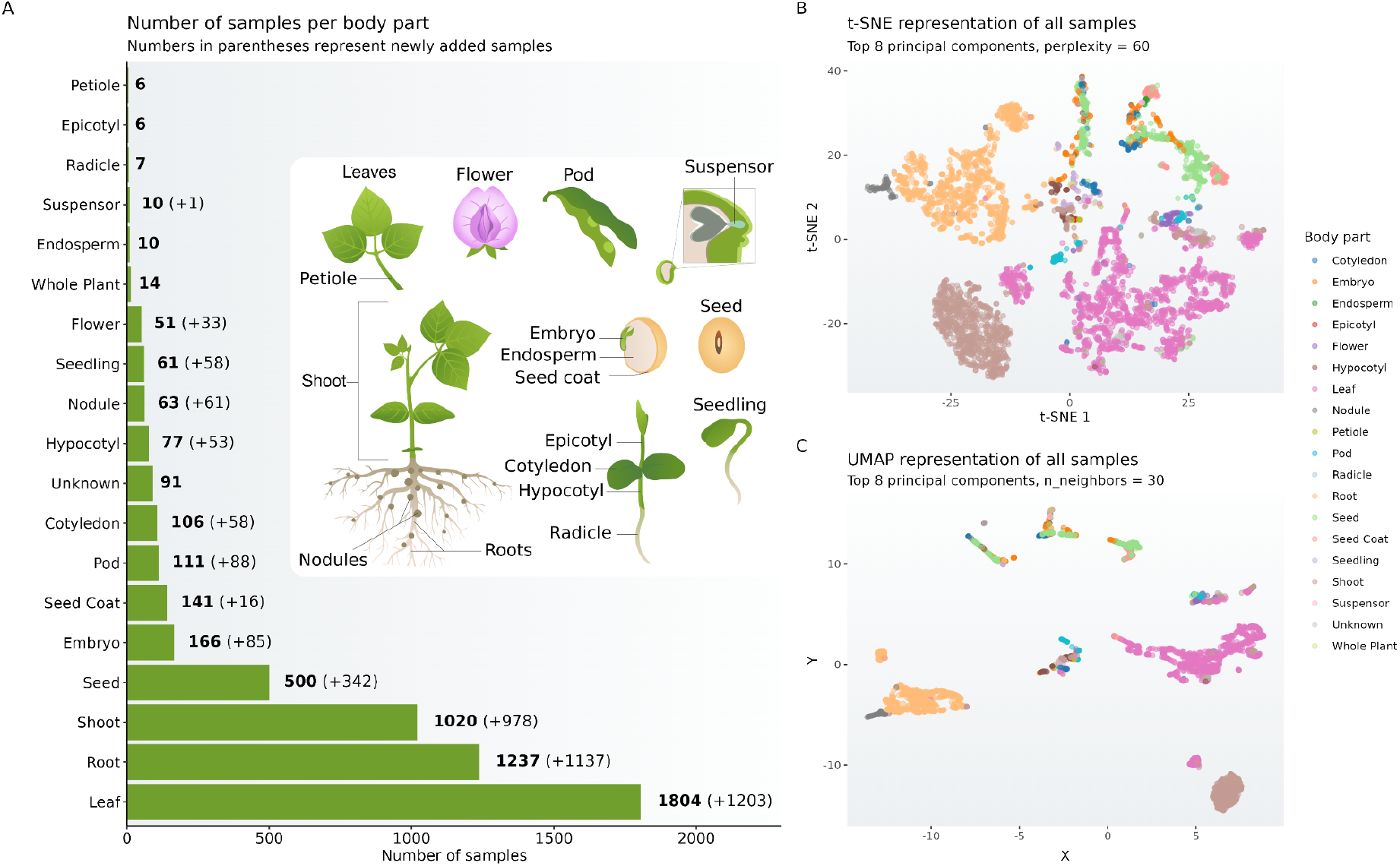
Frequency of samples per body part and dimensionality reduction. **A**. Number of samples per body part. Numbers in bold next to bars represent the number of samples, and numbers in parentheses represent the increase in number of samples relative to the previous version of the Soybean Expression Atlas. **B**. t-SNE representation of samples. t-SNE coordinates were calculated from the top 8 principal components, with *perplexity=60*. **C**. UMAP representation of samples. UMAP coordinates were calculated from the top 8 principal components, with *n_neighbors=30*. Principal components used in the t-SNE and UMAP representations were obtained from a matrix of log-transformed count values for the top 5000 genes with the highest biological components (see Materials and Methods for details).

To have a more detailed view of the quality metrics, we explored the distributions of sequence quality and mapping statistics. We observed that only a small fraction of the samples had mean read lengths <40, and nearly all of the samples had Q20 rates >80%, indicating that the removal of 158 samples in the first quality checkpoint was mainly due to reads being too short (Figure 1C and 1D). The distribution of mapping rates revealed that most of the samples had high mapping rates, but two small peaks can be seen around rates of 35% and 0% (Figure 1E). These samples with low mapping rates represent experiments that: i. aim at quantifying abundances of non-coding regions (e.g., small RNAs) or; ii. inoculate soybean tissues with microorganisms and aim at quantifying bacterial transcript abundances within soybean tissues. The median and mean number of reads per sample were 30 and 39 million reads (Figure 1F). The summary statistics indicate that soybean RNA-seq data in public databases overall have good quality.

### Newly deposited BioSamples reveal the state of the soybean genomics field

We calculated the frequency of samples for each body part and observed that leaf, root, shoot, seed, and embryo had the largest number of samples, with 1804, 1237, 1020, 500, and 166 samples, respectively (Figure 2A). Compared to the previous atlas version (Machado *et al*., 2020), there was a 12-fold increase in the number of root samples, which moved from the fourth to the second position in the ranking by number of samples. When considering only newly deposited samples, the number of root samples was close to the number of leaf samples (*n* = 1137 and 1203, respectively), suggesting that root might overtake leaf in the near future (Figure 2A). Importantly, body parts that had a negligible sample size in the previous version of the atlas, such as nodule and flower (*n =* 2 and 18, respectively), are currently represented by significantly larger sample sizes (*n* = 63 and 51, respectively), which allows more meaningful and reliable conclusions from such data.

A considerable amount of newly deposited root samples represent microbial inoculation experiments with both pathogenic and plant growth-promoting microorganisms, revealing increasing global efforts by the soybean genomics community to understand soybean-microorganism associations (Almeida-Silva *et al*., 2021). Likewise, newly generated samples comprising seed parts (*i*.*e*., seed, embryo, and cotyledon) mostly aim at understanding oil biosynthesis at the transcriptional level, suggesting that this field of research has driven much attention in recent years. Collectively, our findings demonstrate that newly deposited RNA-seq samples can be used as proxies to summarize the current state of the soybean genomics field.

### Transcript abundance differences between body parts explain sample clustering

We performed dimensionality reduction with t-SNE and UMAP using the top 8 principal components to explore global patterns in transcript abundance similarities (Figure 2B and 2C). We found that samples clustered well by body part in both the t-SNE and UMAP representations, but the UMAP representation led to denser clusters with larger distances between them (Figure 2B and 2C). The separation in clusters was unclear for some body parts, which is due to their close anatomical proximity and similarity. For instance, some embryo samples were nearly indistinguishable from endosperm samples, and some nodule samples were very close to root samples (Figure 2B). Likewise, seed samples were highly heterogeneous, with some samples closer to embryo, and other samples closer to seed coat (Figure 2B). As seeds are complex organs that comprise multiple tissues, their larger spread in t-SNE and UMAP representations is unsurprising.

The UMAP and t-SNE representations are consistent with the expected clustering patterns, with samples grouping by body part. Although some authors argue that dimensionality reduction with UMAP better preserves the global structure of the data as compared to t-SNE (Becht *et al*., 2019), others have demonstrated that these methods perform equally well, with differences in global structure attributed to initialization choices (Kobak and Linderman, 2021). However, UMAP has been shown to produce denser and more compact clusters as compared to t-SNE, with more space between clusters (Kobak and Linderman, 2021), which is indeed what we observed (Figure 2B and 2C). Despite the differences between methods, both the t-SNE and UMAP representations reveal that samples cluster by body part.

### Geographic patterns in data deposits and the evolution of sequencing technologies

We calculated the number of samples per country to explore geographical patterns in soybean RNA-seq data generation. We observed that world leaders in soybean production (*i*.*e*., US, China, Brazil, Canada, and India) are also world leaders in soybean RNA-seq data generation (Figure 3A and 3B) (http://soystats.com). Strikingly, the number of samples deposited by institutions in the US and China (*n* = 3023 and 1405) is much larger than the number of samples contributed by other countries. For instance, the number of samples contributed by China is greater than the number of samples contributed by all other countries combined (excluding the US), and the number of samples contributed by the US is twice as large as that of China.

**Fig. 3.**
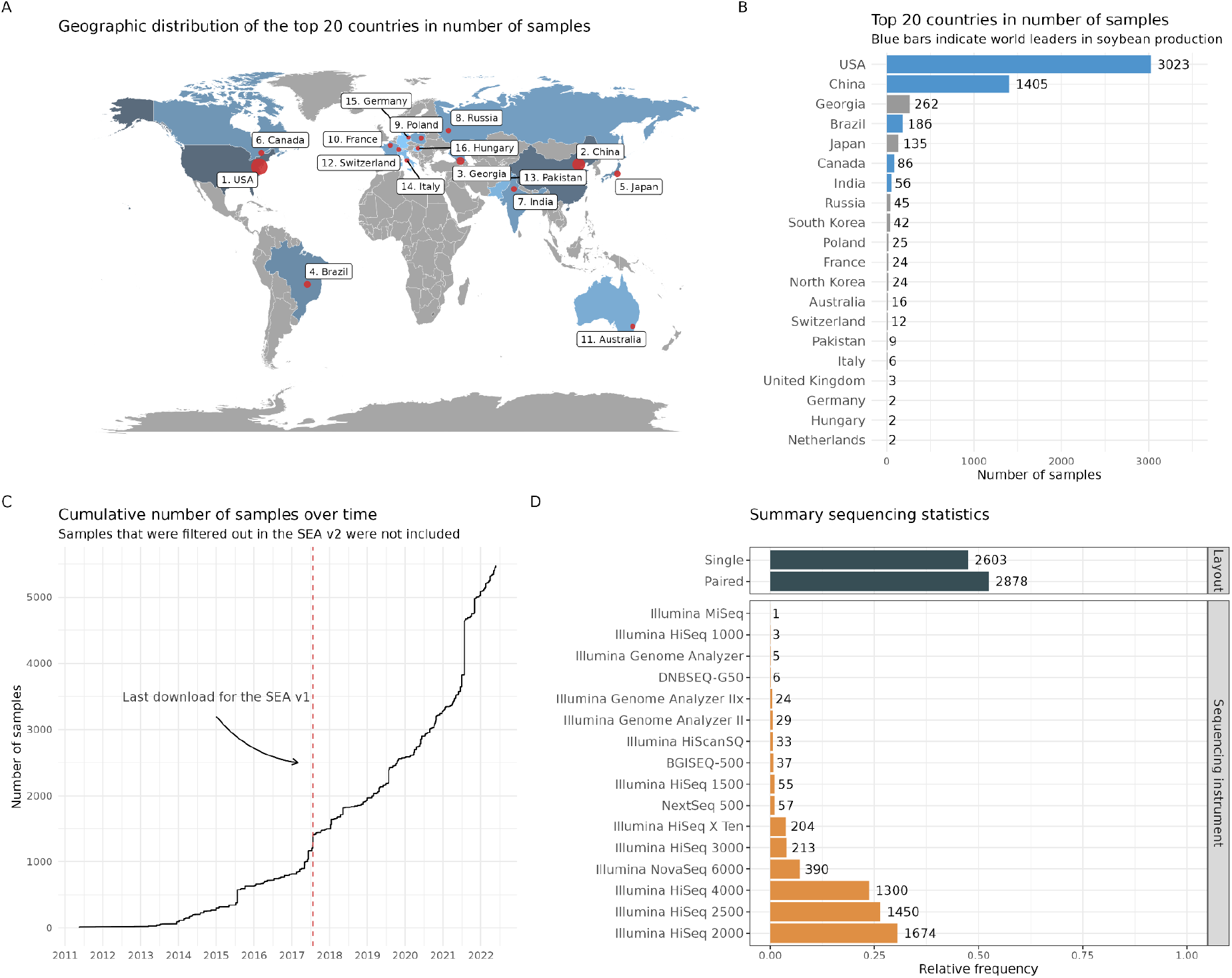
Global distribution of sample deposits and summary sequencing statistics. **A**. Geographic distribution of the top 20 countries in number of deposited samples. **B**. Bar plot of the top 20 countries in number of deposited samples. Blue bars represent countries that are world leaders in soybean production. **C**. Cumulative number of samples over time. Samples that failed the quality checkpoints were not included in the visualization. **D**. Summary statistics on library layout and sequencing instruments.

Further, we explored summary sequencing statistics to better understand how such data were generated. Most of the samples were generated with paired-end sequencing (*n* = 2878), and Illumina HiSeq 2000, HiSeq 2500, and HiSeq 4000 were the most commonly used sequencing instruments (*n* = 1674, 1450, and 1300, respectively) (Figure 3D). Sample deposits over time show that new samples accumulate exponentially (Figure 3C), which highlights the challenges of maintaining a database such as the Soybean Expression Atlas and the need for stable funding. The frequency rankings by body part and sequencing instrument have mostly remained the same over time, but the time-series visualizations reveal when changes occurred, such as root samples overtaking seed samples in 2018 (Supplementary Figure S2A), the sharp increase in number of samples generated with Illumina NovaSeq 6000 since 2021 (Supplementary Figure S2C), and paired-end sequencing overtaking single-end sequencing in 2019 (Supplementary Figure S2B).

### A new web interface to keep up with the increasingly complex data sets in databases

We developed a new web interface, available at https://soyatlas.venanciogroup.uenf.br/, to allow easier and more meaningful search, exploration, and download of the expression data in the Soybean Expression Atlas v2. To preserve the structure of the previous version, the new interface comprises 4 pages: *Global Expression Viewer, Search gene list, Download by body part*, and *Download by project*. All pages except *Download by body part* have been updated to include features that enhance user experience and interpretation of results. The web interface allows exploration and download of gene-level transcript abundances, but transcript-level abundances are also available as *SummarizedExperiment* objects in the FigShare repository associated with this publication (Almeida-Silva and Venancio, 2023b).

The *Global Expression Viewer* allows users to enter a gene ID and visualize mean and median expression profiles per body part. However, as the Soybean Expression Atlas v2 is highly heterogeneous, with several different developmental stages and treatments for the same tissue, we now allow users to visualize the expression profile of a gene in each individual sample by using t-SNE coordinates (Figure 4A). This new feature can help users detect if the expression of the input gene in a plant part is consistent across all samples or if there are condition specific-variations (*e*.*g*., a gene is highly expressed in water stress-induced roots, but not in control conditions).

**Fig. 4.**
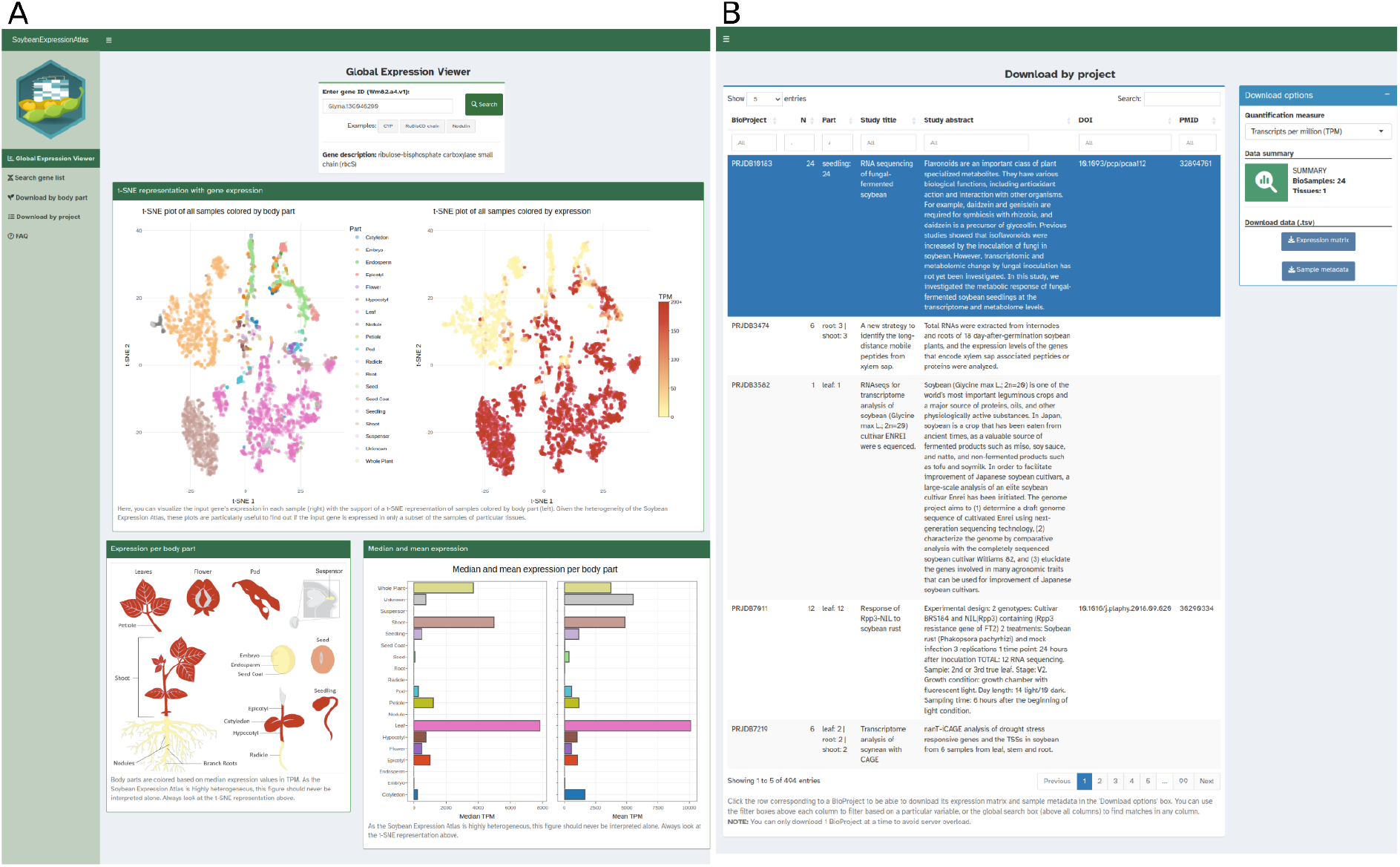
Example features of the new web application. **A**. *Global Expression Viewer* page highlighting the gene-level transcript abundance profiles of a ribulose-1,5-bisphosphate carboxylase/oxygenase (RuBisCO) subunit. Upon search, users can visualize transcript abundances based on t-SNE coordinates, a map of plant body parts, and bar plots of mean and median abundances. **B**. *Download by project* page. After selecting a given BioProject in the interactive table (highlighted in blue), users can download gene expression matrices and/or sample metadata as tab-separated files.

The *Search gene list* page allows users to enter a gene list (up to 200 genes) and visualize their expression profiles (in TPM or bias-corrected counts) in selected body parts as a heatmap. Users can also download the gene expression matrix used to create the heatmap and sample metadata as tab-separated files. In the latest interface, we updated this page to include body part annotations and hierarchical clustering of genes in the heatmap, which facilitates interpretation.

Finally, the *Download by project* page allows users to download the gene expression matrix (in TPM or bias-corrected counts) for a particular BioProject. Previously, users could search projects by DOI or PubMed ID, which restricted download to only projects that have associated publications. In the latest interface, we included an interactive table of BioProject metadata with column filters and a search bar that facilitates the search for projects of interest (Figure 4B). For instance, users can find and download projects containing more than *N* samples, projects that contain a body part of interest, or projects that contain a user-defined text string in their titles or abstracts (e.g., “water stress”, “seed oil”, “disease”, etc).

## Conclusions

In this work, we present an updated version of the Soybean Expression Atlas, a gene expression database of 5481 high-quality RNA-seq samples publicly available on the NCBI’s SRA processed with a standardized pipeline. Gene-level transcript abundances in TPM and bias-corrected counts can be easily explored and downloaded from the web application, which will keep empowering soybean researchers towards discoveries. Transcript-level abundances and equivalence classes are also available to allow differential transcript usage analyses. The easily accessible expression data in our database can be used for various purposes, such as differential expression analyses, functional studies, and identification of genes involved in important traits.

## Acknowledgements

FA-S acknowledges funding from Ghent University (Methusalem funding, BOF.MET.2021.0005.01). FP-S and TMV acknowledge funding from Fundação Carlos Chagas Filho de Amparo à Pesquisa do Estado do Rio de Janeiro (FAPERJ; grant E-26/203.014/2018), Coordenação de Aperfeiçoamento de Pessoal de Nível Superior—Brasil (CAPES; Finance Code 001), and Conselho Nacional de Desenvolvimento Científico e Tecnológico. The funding agencies had no role in the design of the study and collection, analysis, and interpretation of data and in writing.

## Data availability

All data and code used in this paper are available in a GitHub repository (https://github.com/almeidasilvaf/SEA_paper) to ensure full reproducibility.

