## Supplementary Figures for "The Soybean Expression Atlas v2: a comprehensive database of over 5000 RNA-seq samples"

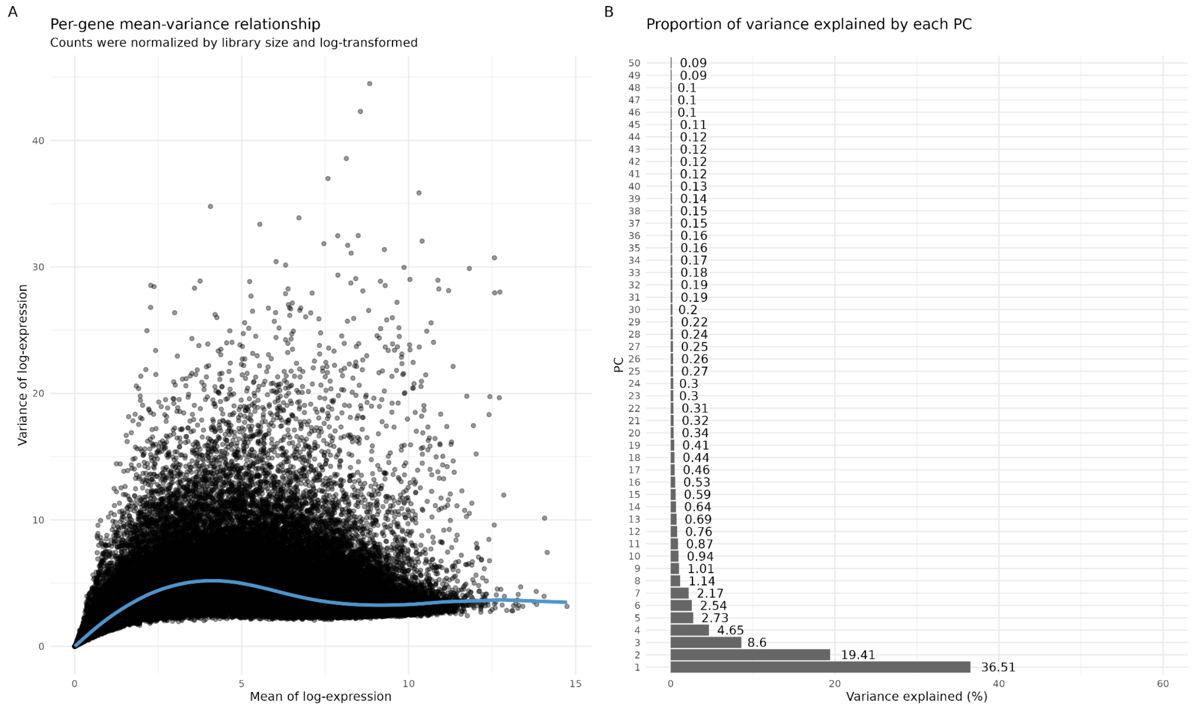


**Fig. 1. Feature selection for dimensionality reduction. A.** Per-gene mean-variance relationship obtained with the *modelGeneVar()* function from the scran package. The model was fitted on a matrix of log-transformed counts normalized by library size. **B.** Proportion of variance explained by each principal component. The top 8 principal components were selected for dimensionality reduction, as they explain most of the variation.

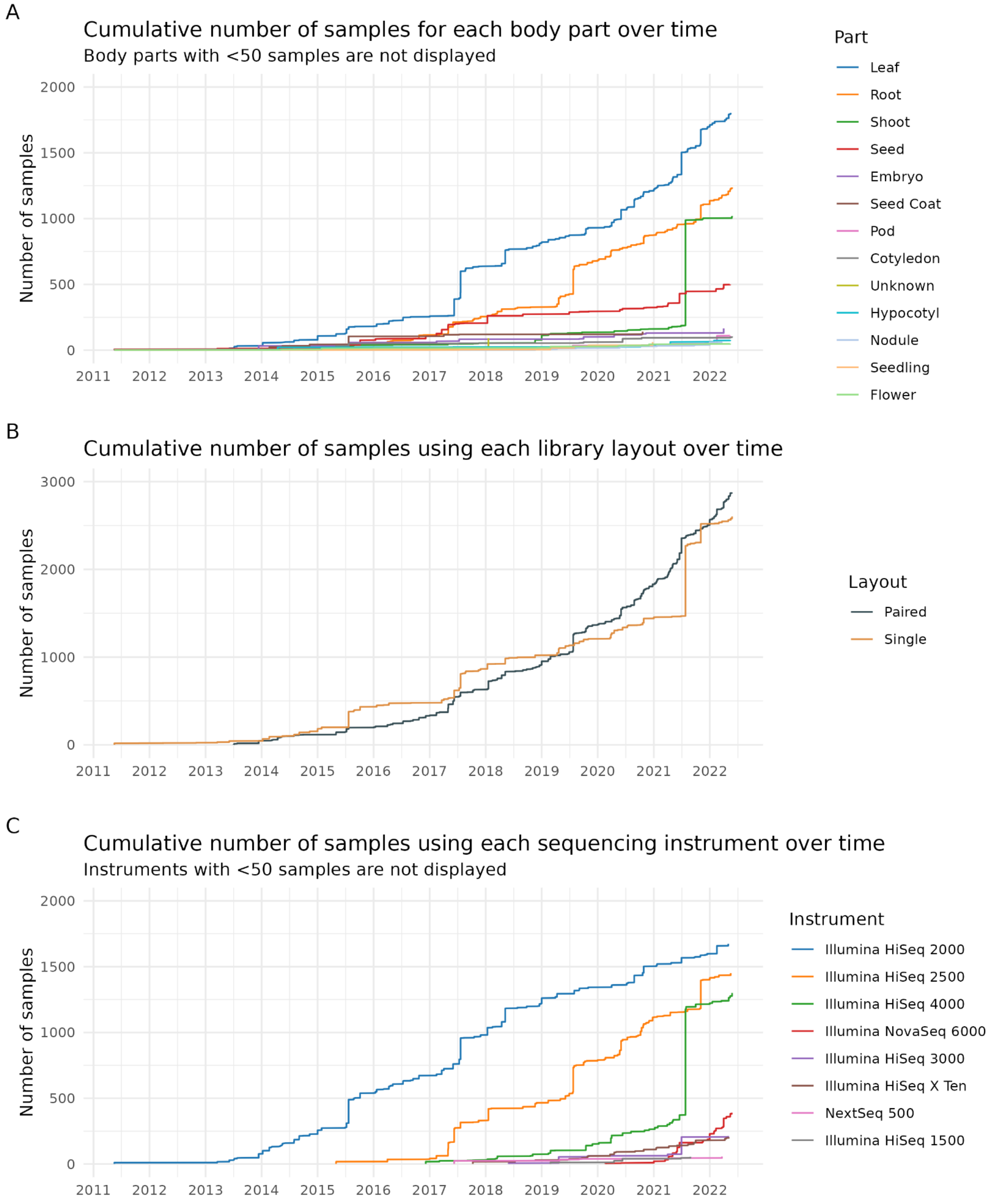


**Fig. 2. Summary statistics in a time series. A.** Cumulative number of samples per body part over time. Body parts represented by less than 50 samples were not included in the visualization. **B.** Cumulative number of samples per library layout over time. **C.** Cumulative number of samples per sequencing instrument over time. Instruments represented by less than 50 samples were not included in the visualization.
